# Induction of muscle stem cell quiescence by the secreted niche factor Oncostatin M

**DOI:** 10.1101/303586

**Authors:** Srinath C. Sampath, Srihari C. Sampath, Andrew T.V. Ho, Stéphane Y. Corbel, Joshua D. Millstone, John Lamb, John Walker, Bernd Kinzel, Christian Schmedt, Helen M. Blau

## Abstract

The balance between stem cell quiescence and proliferation in skeletal muscle is tightly controlled, but perturbed in a variety of disease states. Despite progress in identifying activators of stem cell proliferation, the niche factor(s) responsible for quiescence induction remain unclear. Here we report an *in vivo* imaging-based screen which identifies Oncostatin M (OSM), a member of the interleukin-6 family of cytokines, as a potent inducer of muscle stem cell (MuSC, satellite cell) quiescence. OSM is produced by muscle fibers, induces reversible MuSC cell cycle exit, and maintains stem cell regenerative capacity as judged by serial transplantation. Conditional OSM receptor deletion in satellite cells leads to stem cell depletion and impaired regeneration following injury. These results identify Oncostatin M as a secreted niche factor responsible for quiescence induction, and for the first time establish a direct connection between induction of quiescence, stemness, and transplantation potential in solid organ stem cells.

## Introduction

Stem cells respond to tissue-specific activating signals by proliferating and giving rise to both committed progenitors as well as quiescent daughter cells. Muscle repair is mediated by resident muscle stem cells (MuSC; also known as satellite cells). In response to myofiber damage, satellite cells break from quiescence and give rise to committed progenitors known as myoblasts, which can fuse both with each other as well as with the damaged myofibers. Importantly, a subset of activated satellite cells are then re-selected into the quiescent stem cell niche in order to maintain the stem cell pool^1^. This process is highly regulated, involving not only cell autonomous signaling pathways, but also extensive regulation by tissue resident stromal cells and invading inflammatory cell populations^2^. These regulatory pathways are critical, as defects in stem cell quiescence, activation, or self-renewal have been implicated in a variety of disease states including aging-associated sarcopenia, muscular dystrophy, and cancer cachexia^3, 4, 5, 6^.

Significant progress has been made toward elucidating the complex extracellular cues arising from the circulation and microenvironment that govern the behavior of tissue resident MuSC. Among the known soluble growth factor/receptor pathways, the HGF pathway has been demonstrated to control ‘alerting’ of satellite cells in response to remote injury^7^, inducing a state of metabolic activation and priming for cell cycle reentry. FGF likewise regulates satellite cell activation^8^, while a downstream negative regulator of FGF receptor tyrosine kinase signaling, Sprouty1, plays an important role in the acquisition of quiescence of activated satellite cells ^9^. Growth factor signaling eventually impinges on the p38α/β MAPK pathway, which both genetic and pharmacologic data suggests functions as an important regulator of stemness and proliferation^6, 10^. Additional levels of regulation are imposed via Notch/Delta-dependent regulation of self-renewal^11, 12, 13^, Wnt-dependent control of myogenic fate ^14^, and asymmetric division associated with differential Pax7 expression^15^.

In addition to the growth factor pathways described above, previous work has demonstrated the importance of circulating cytokines in regulating MuSC and myofiber function. The IL-6 pathway has been a particular focus given the upregulation of this pleiotropic cytokine following exercise^16^, as well as the aberrant regulation of the downstream JAK-STAT signaling pathway during aging and in other disease states^17^. Studies with *IL-6* deficient animals indicated a role for this cytokine in satellite cell and myoblast proliferation during experimentally-induced hypertrophy^18^. The IL-6 family member Leukemia Inhibitory Factor (LIF) has likewise been demonstrated to induce proliferation of human and mouse myoblasts^19^. Indeed, it has recently been reported that inhibition of STAT3-dependent signaling, which is activated by both LIF and IL-6, can rescue age-associated proliferative defects in mouse satellite cells^20^ and promote satellite cell expansion^21^. Interpretation of the results of genetic and pharmacologic STAT3 inhibition is complicated, however, by the fact that this effector can be activated by numerous upstream signals, including both IL-6 and other cytokines^22^.

Despite this progress, the precise signals governing the choice between quiescence and proliferation remain poorly understood, and this constitutes a major barrier to reversing the regenerative block seen in a variety of disease states. In particular, the identities of the factors regulating quiescence of adult stem cells have remained elusive. While bioinformatic analyses have begun to address this issue by identifying genes enriched in quiescent stem cells^23, 24^, the functional significance of most of these putative regulators has not been established. To address this deficiency, we have undertaken a systematic screen for secreted and transmembrane protein regulators of stem cell proliferation and engraftment. Employing an unbiased *in vitro/in vivo* imaging based screening strategy, we identify Oncostatin M, a member of the IL-6 family of cytokines, as a novel regulator of stem cell quiescence, engraftment, and muscle regeneration.

## Results

### A secretomic screen for muscle stem cell regulators

We hypothesized that a secreted regulator of stem cell quiescence would be capable of suppressing MuSC activation and proliferation *in vitro*, while conversely maintaining the ability of these cells to engraft following transplantation *in vivo*. To identify such regulators, we designed and carried out a protein library screen on purified MuSCs, using a combination of *in vitro* and *in vivo* imaging to distinguish between effects on stem cell proliferation and engraftment (Fig. 1a). Primary muscle stem cells isolated from *luciferase* transgenic mice^25, 26^ were cultured on uncoated plastic plates in the presence of pools of proteins derived from the GNF Secretomics library, which contains purified versions of known or predicted mammalian secreted proteins, as well as the extracellular domains of known or predicted single-pass transmembrane proteins^27^. No additional growth factors (e.g. FGF) were added. *In vitro* bioluminescence imaging (BLI) was used to assess stem cell proliferation (Supplementary Fig. 1a, top), followed by cell harvesting and transplantation into the *tibialis anterior* (TA) muscle of hindlimb-irradiated recipients to assess engraftment, a sensitive measure of regeneration (Supplementary Fig. 1a, bottom). Serial *in vivo* imaging was performed over a timecourse to quantify engraftment activity of the transplanted cells, as indicated by a progressive increase in light output, a feature of stem cells but not committed progenitors^25^. A comparison was made to both untreated cells, as well to known inducers of stem cell proliferation, FGF and the p38 MAPK inhibitor, SB202^6, 10^. Protein pools demonstrating increased engraftment on serial *in vivo* imaging were iteratively deconvolved through rounds of subpool rescreening to identify the single protein responsible for the engraftment activity (Fig. 1b).

**Figure 1.**
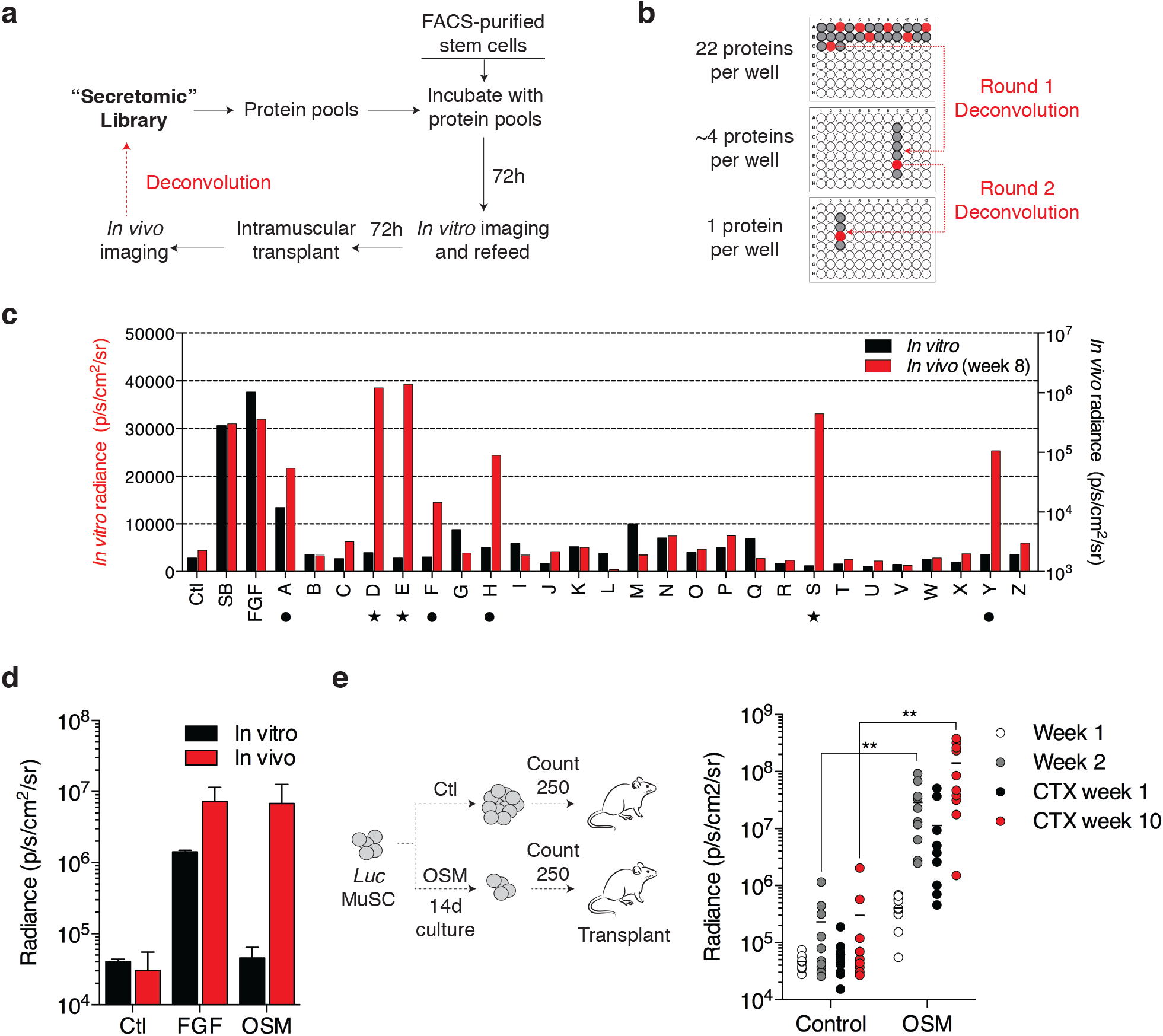
Identification of Oncostatin M as a potent inducer of muscle stem cell engraftment. a. Schematic of combined *in vitro/in vivo* bioluminescence imaging (BLI) screening strategy.
b. Schematic of deconvolution for positive protein pools. Pools of 22 proteins scoring for engraftment in the initial round of screening were iteratively deconvolved into pools of 4 proteins, and then individual proteins.
c. Results of multiplexed *in vitro/in vivo* protein screening. Black, results of *in vitro* BLI; red, results of *in vivo* BLI. Ctl: control buffer; SB: SB202190; FGF: recombinant human basic Fibroblast Growth Factor. Symbols below the axis mark pools containing OSM (star), and all other hits (circle; includes FGF, TNFR, and incompletely deconvolved pools).
d. Representative BLI results following treatment of 800 FACS-sorted MuSC with either FGF or Oncostatin M (OSM; 100 ng/mL) for 6 days *in vitro*, followed by intramuscular transplantation (3 8-12 week old animals each) and *in vivo* BLI measurement 6 days (open circles) or 2 weeks (shaded circles) after transplant.
e. Schematic (left) demonstrating transplantation of 250 counted progeny of CD11b^−^Sca1^−^CD31^−^CD45^−^CD34^+^α7 integrin^+^ stem cells following long-term 2 week culture with OSM control conditions (FGF). Serial BLI following transplantation of counted cells (right). Each point indicates an individual hindlimb that received a 250 cell transplant, n=10 per condition. BLI is shown for weeks 1 (white) and 2 (light grey) pre-CTX, and weeks 1 (dark grey) and 10 (red) post-CTX. Double asterisks indicate p<0.01 in comparison with control condition at the same timepoint.

Pooled screening of ~600 predominantly mouse proteins yielded 5 hits in two phenotypic classes. The first class was exemplified by a pool driving both *in vitro* proliferation as well as *in vivo* engraftment (pool A; Fig. 1c, Supplementary Fig. 1b). Iterative deconvolution identified the responsible protein as FGF2, a known mitogenic factor for MuSC^28^, providing strong validation of the assay. The second phenotypic class exemplified the hypothesized behavior for an inducer of reversible stem cell quiescence, driving little *in vitro* stem cell proliferation but robust *in vivo* engraftment. Deconvolution of one such pool (pool Y; Fig. 1c, Supplementary Fig. 1b) identified the extracellular domain of the TNFα receptor (Supplementary Fig. 1c), which acts as a receptor decoy for TNFα and which is consistent with the known pro-differentiation function of TNFα on satellite cells^29^. Pools F and H were not able to be completely deconvolved, possibly due to partial contribution of multiple protein factors. Strikingly, deconvolution of the remaining pools in this phenotypic class identified the same protein in three differentially epitope-tagged forms, Oncostatin M (OSM; pools D, E, and S; Fig. 1c, star; Supplementary Fig. 1b). No engraftment-promoting activity was seen in subpools lacking Oncostatin M (Supplementary Fig. 1d,e). Recombinant OSM obtained from commercial sources demonstrated similar engraftment-inducing activity (Fig. 1d). Histological analysis of engrafted recipient mice confirmed the presence of donor-derived myofibers (Supplementary Fig. 2a), in keeping with the good correlation between BLI and histology seen previously^6, 25^.

Single cell RT-PCR analysis of purified MuSC demonstrated expression of the Oncostatin M receptor (OSMRβ) within a subset comprising ~20% of Pax7^+^ cells (Supplementary Fig. 2b), which may reflect either true heterogeneity, or limited sensitivity of the single cell detection method. To exclude effects on proliferation as a confounding variable, stem cell engraftment was next determined on a per-cell basis. Satellite cells were cultured for 2 weeks either with OSM or in standard medium containing FGF, and intramuscular transplant of 250 counted cells was then performed (Fig. 1e, left). Serial *in vivo* imaging revealed progressive increase of light output, indicative of engraftment, and injury of the stem cell-engrafted muscle with cardiotoxin led to a drop in BLI signal followed by a robust rebound, confirming the presence of functional satellite cells (Fig. 1e, right). BLI signal was significantly higher in muscle receiving OSM treated MuSC, consistent with increased per-cell engraftment and regeneration.

### Regulation of muscle stem cell quiescence by OSM

To further investigate the anti-proliferative effect of OSM, we performed serial *in vitro* BLI. This demonstrated inhibition of stem cell proliferation at all timepoints (Fig. 2a, Supplementary Fig. 3a), which was attributable to decreased S-phase entry (Fig. 2b). Proliferation of stem cells activated by culture on muscle fibers *ex vivo* was also strongly inhibited (Supplementary Fig. 3b,c), and OSM treatment of primary mouse myoblasts likewise decreased levels of the M-phase marker phospho-H3S10 (Fig. 2c). Previous studies have demonstrated a critical role of the cyclin-dependent kinase inhibitor p27/Kip1 in quiescence of stem cells both in muscle as well as other lineages^30, 31^. We also observed a significant decline in *p27* transcript in cultured MuSC, and moreover found that OSM treatment maintained *p27* expression at levels seen in freshly isolated, non-cultured stem cells (Supplementary Fig. 4a), suggesting that OSM might directly induce quiescence. To explore this possibility, we compared the global transcriptional profile of freshly isolated MuSCs to MuSCs cultured for 3 days with or without OSM, as well as to prior transcriptional studies of muscle stem cell quiescence^23, 24^ The published analyses were used to extract a transcriptional signature of quiescence regulated genes. A comparison of this ‘quiescence signature’ with the transcriptional response to OSM revealed highly significant enrichment (Fig. 2d), which was independently confirmed by qPCR analysis (Supplementary Fig. 4b). Over half of the signature genes were present within the OSM-regulated set (Supplementary Data 1), consistent with induction of a global transcriptional quiescence program.

**Figure 2.**
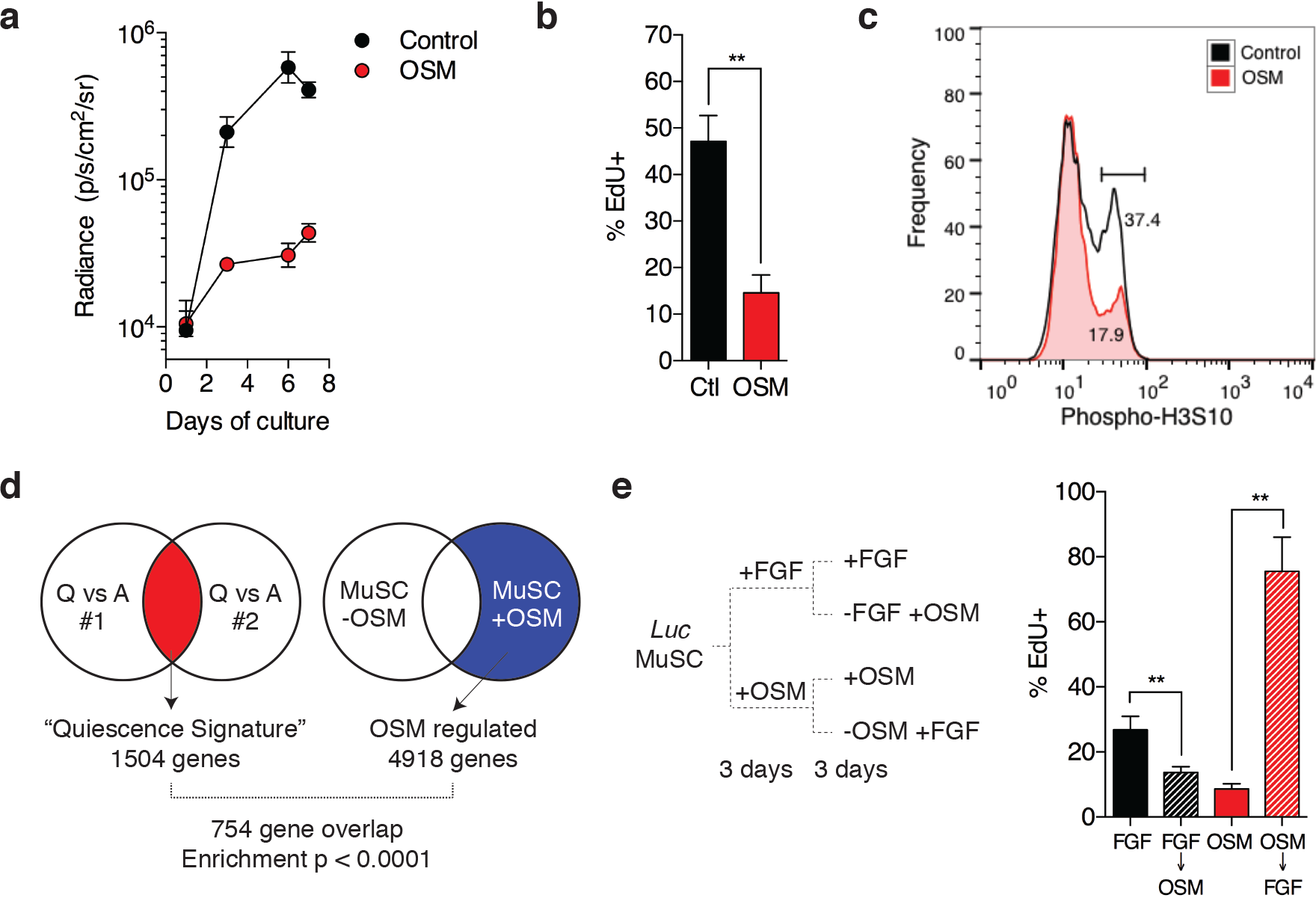
OSM induces reversible cell cycle exit and a molecular signature of quiescence. a. FACS-sorted muscle stem cells purified from *luciferase* transgenic mice were cultured for 7 days at 2500 sorted counts per well (~31 cells/mm^2^) in the presence (closed circle) or absence (open circles) of OSM, as well as low dose FGF (0.5 ng/ml). Serial *in vitro* BLI was performed at the indicated timepoints. Mean +/- SD is shown, n=4 replicates.
b. FACS-sorted MuSC cultured for 6 days as in (A) were pulsed with EdU for the last 12 hr of culture. Mean +/- SD is shown, n=4 replicates.
c. Induction of cell cycle exit in myoblasts in response to OSM. Primary myoblasts were cultured for 3 days in growth medium with (red) out without (grey) added OSM. Cells were fixed, intracellular staining performed for mitotic H3S10 phosphorylation, and analyzed by flow cytometry.
d. Diagram demonstrating overlap between a bioinformatically-derived gene expression “quiescence signature” for satellite cells, and genes regulated during OSM-treatment of muscle stem cells *in vitro*. OSM regulation data was derived mean of 2 biological replicates.
e. Schematic of switch experiment to demonstrate reversibility of OSM-induced cell cycle exit (left). FACS-sorted MuSC were cultured for 6 days under the indicated conditions, and then pulsed with EdU for the final 12 hr of culture (right). Mean +/- SD is shown, n=4 replicates; greater than 300 cells were counted per replicate.

Reversibility is the hallmark feature of quiescence, and we therefore tested whether OSM-induced cell cycle exit was reversible by culturing satellite cells in either activating (-OSM) or non-activating (+OSM) conditions for 3 days, after which the medium conditions were switched and EdU pulse labeling performed (Fig. 2e, left). Non-cycling OSM-treated cells robustly re-entered cell cycle following stimulation, as measured by both EdU incorporation (Fig. 2e, right) and cell proliferation (Supplementary Fig. 4c); This anti-proliferative effect on satellite cells was dependent on MEK activity (Supplementary Fig. 4d), consistent with prior reports of OSM activity in other cell types^32, 33^.

### OSM treatment maintains stemness in vitro

Although both satellite cells and their committed progeny (myoblasts) express Pax7 and can contribute to myofiber formation^34^, only satellite cells are capable of mediating serial regeneration following injury^25^. The ability to reisolate and retransplant engrafted satellite cells is thus a strict test of stem cell identity, and we therefore tested whether OSM-treated MuSC fulfill these criteria. We employed a newly generated reporter allele in which an Internal Ribosome Entry Site and Green Fluorescent Protein (IRES-GFP) are inserted into the 3’ untranslated region of the *Pax7* locus (Pax7^GFP^; Supplementary Fig. 5a). This allele specifically labels CD34^+^/α7 integrin^+^ satellite cells in skeletal muscle (Supplementary Fig. 5b). MuSC from Pax7^GFP^ mice were cultured in the presence or absence of OSM, and intramuscular transplant of equal numbers of treated stem cells was performed. Despite high initial engraftment of cells cultured either with or without OSM (Fig. 1d), Pax7^GFP^-expressing MuSC could only be reisolated from mice engrafted with OSM-treated cells (Fig. 3a). To establish that the engrafted cells represent *bona fide* functional stem cells, Pax7^GFP+^ cells were re-isolated from primary recipients of OSM-treated MuSC, and 50 counted cells were transferred into the hindlimbs of irradiated secondary recipients. BLI demonstrated engraftment in ~40% of recipients, and cardiotoxin injury induced initial loss followed by rebound of BLI signal, a pattern characteristic of stem cell-mediated regeneration (Fig. 3b). Previous studies have demonstrated that such increased serial transplantation ability corresponds with higher Pax7 levels^15^, and we likewise found that OSM treatment not only induced *Pax7* (Fig. 3c) but also reciprocally decreased expression of the differentiation marker *myogenin* (Supplementary Fig. 6a). This effect was unique amongst the related IL-6 family of cytokines (Fig. 3d)^20, 21^. Intracellular staining and flow cytometry in primary myoblasts demonstrated that OSM preferentially activated STAT3 rather than STAT5 (Fig. 3e). Analysis of the *Pax7* locus revealed a previously uncharacterized binding site for STAT3, and chromatin immunoprecipitation (ChIP) from primary myoblasts confirmed direct occupancy of this site by STAT3 in response to OSM treatment (Fig. 3f). Taken together, these data reveal a MEK and STAT3-mediated response to OSM in satellite cells, leading to quiescence induction as well as maintenance of stem cell identity and function.

**Figure 3.**
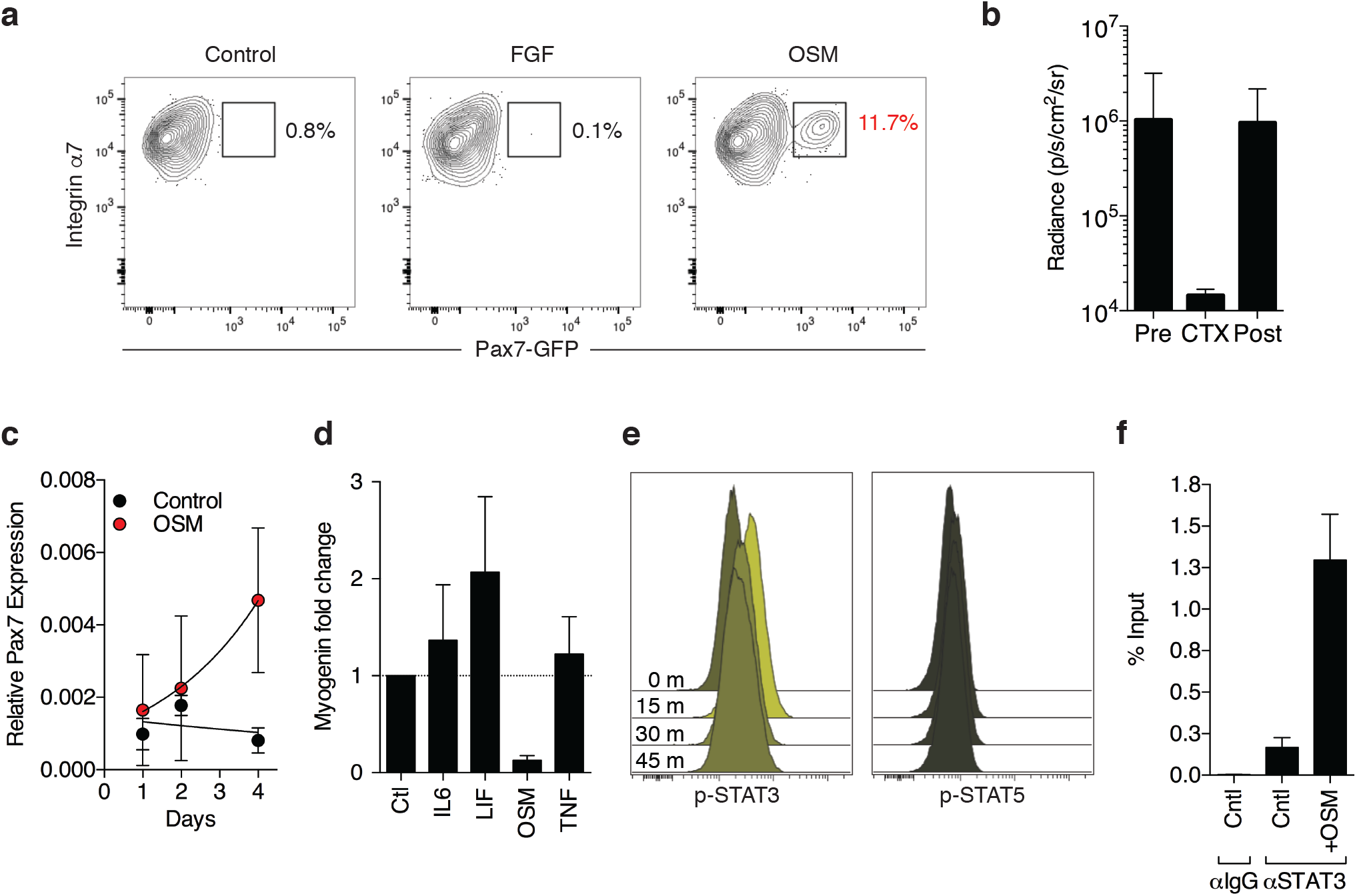
OSM treatment enhances stem cell serial transplantation and inhibits differentiation. a. FACS analysis of stem cells reisolated from primary transplant recipients receiving either control (left), FGF (middle), or OSM-treated (right) stem cells. Only the live, CD11b^−^Sca1^−^CD31^−^CD45^−^CD34^+^α7 integrin^+^ stem cell population is shown.
b. MuSCs (CD11b^−^Sca1^−^CD31^−^CD45^−^CD34^+^α7 integrin^+^GFP^+^) were reisolated from OSM primary transplant mice as described in (A), transplanted equally into 5 limbs (secondary transplant recipients; 50 MuSC/limb; 8-12 week old recipients), and monitored by serial BLI. At day 25 following transplant, cardiotoxin (CTX) was injected to induce myofiber damage and stimulate stem cell expansion and tissue regeneration. The average radiance at each timepoint is indicated, and demonstrates regenerative capacity (BLI signal loss followed by rebound) of serially transplated OSM-treated MuSC. Percentage of engrafted mice was 40% both pre and post-CTX injury (2/5 total mice). Note that untreated cells give no engraftment, and no MuSC are therefore present for reisolation/transplant.
c. qRT-PCR analysis of *Pax7* expression following 6 day culture of FACS-sorted MuSC in the presence of OSM (gray circles) or buffer control (black). Mean +/- SD is shown, n=3 replicates.
d. qRT-PCR analysis of *myogenin* expression following 6 day culture of FACS-sorted MuSC with the indicated cytokines and growth factors. Expression was normalized to GAPDH expression. Mean +/- SD is shown, n=3 replicates.
e. Timecourse intracellular flow cytometry of primary mouse myoblasts treated for the indicated period of time with OSM, and stained for either phospho-STAT3 (left) or phospho-STAT5 (right). 0 (unstimulated), 15, 30, and 45 minute timepoints are shown following cytokine addition. Frequencies are represented as histograms.
f. ChIP performed using primary myoblasts cultured for 6 days either with or without OSM. Immunoprecipitation was performed with either control IgG or STAT3-specific antibodies, and PCR was performed using primers specific for a predicted STAT3 binding site adjacent to the Pax7 locus.

### Functions of OSM signaling in vivo

To address the endogenous source of OSM in muscle, we assessed protein levels in muscle tissue and found they were highest in the resting state, decreased rapidly following injury, and reappeared beginning at day 4 and robustly by day 10, concomitant with myofiber regeneration (Fig. 4a). This suggests that OSM is produced in a paracrine fashion from the myofiber niche, rather than in an autocrine fashion from satellite cells. Moreover, this timecourse is inconsistent with immune cells, such as macrophages, being the primary source of OSM, as reported by others in the heart^33^. Immunofluorescence staining of OSM in uninjured muscle indeed revealed localization within myofibers of uninjured muscle, both around myonuclei and the fiber periphery (Fig. 4b), consistent with OSM production by myofibers as a secreted niche factor. To investigate the physiological significance of niche-induced OSM signaling, we derived mice deficient for the obligate OSM receptor, OSMRβ (Supplementary Fig. 6b). Intermediate OSMRβ levels were present in heterozygous animals (Supplementary Fig. 6b), consistent with previously reported haploinsufficiency^33^. EdU incorporation was significantly higher in OSMR-deficient satellite cells (Supplementary Fig. 6c), demonstrating that OSM signaling in satellite cells restrains stem cell proliferation. Importantly, OSMRβ was required for the pro-quiescence effect of OSM (Supplementary Fig. 6c), excluding the possibility of additional receptors mediating this signal.

**Figure 4.**
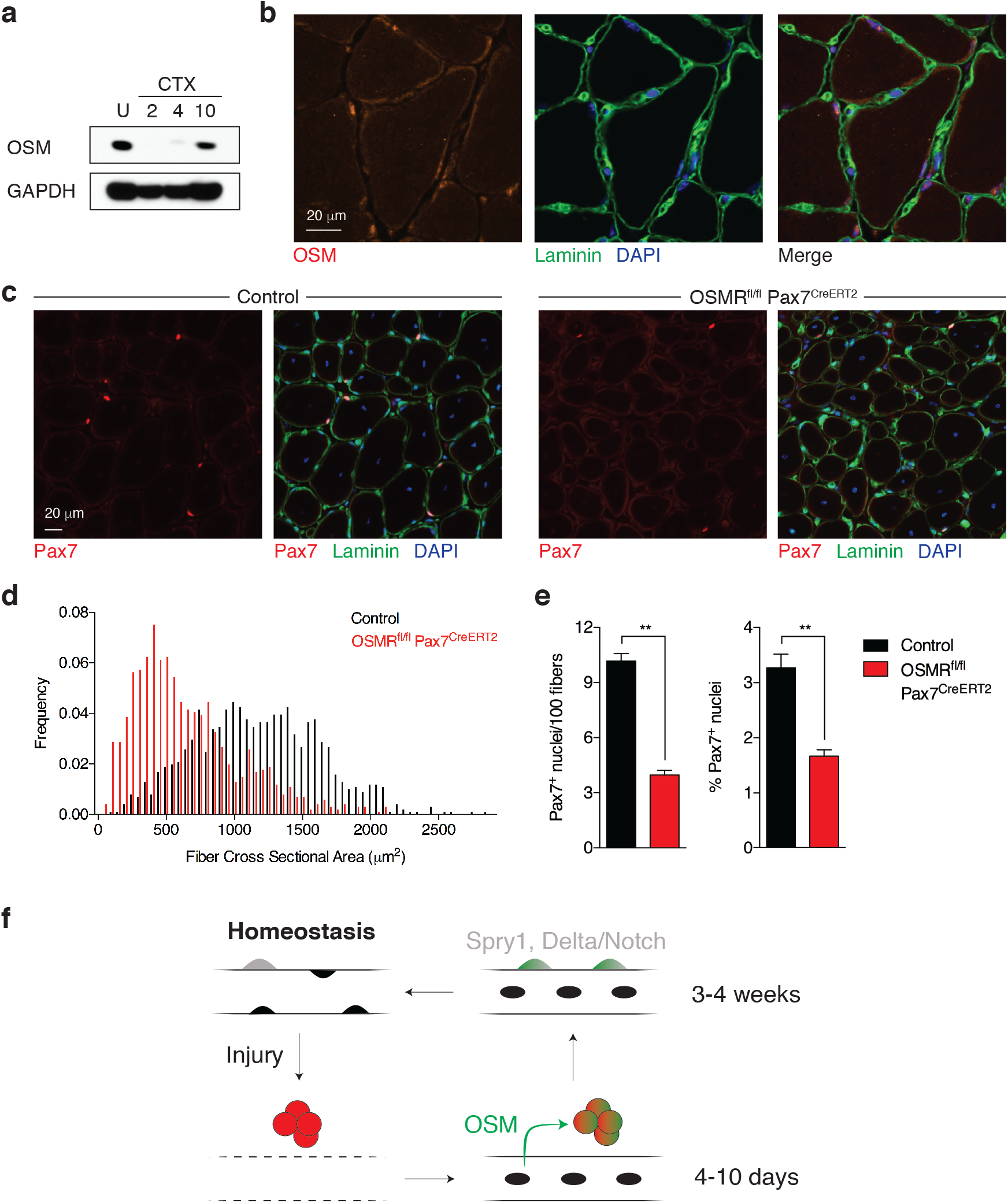
OSM signaling regulates quiescence induction and is required for proper muscle regeneration. a. Western blot of tibialis anterior muscle, either uninjured (U) or at the indicated days following cardiotoxin injury.
b. Immunofluorescence analysis of wild type uninjured TA muscle. Blue: DAPI; Green: Laminin; Red: OSM. Note the expected perinuclear staining pattern of OSM, consistent with synthesis and export through the known perimyonuclear ER/Golgi network^49, 50^.
c. Representative immunofluorescence images from tibialis anterior muscle of control (Osmr^fl/+^) or conditional knockout (Osmr^fl/fl^ Pax7^CreERT2^) animals following deletion and 2 rounds of cardiotoxin injury and regeneration. For each sample, single channel Pax7 staining is shown on the left and a merged 3 channel image is shown on the right. Blue: DAPI; Green: Laminin; Red: Pax7.
d. Cross-sectional area (CSA) distribution of myofibers from TA muscle of control (Osmr^fl/+^) or conditional knockout (Osmr^fl/fl^ Pax7^CreERT2^) animals (3 8-12 week old animals each) following deletion and 2 rounds of cardiotoxin injury and regeneration. >1000 fibers were quantified per genotype.
e. Quantitation of Pax7^+^ stem cells in serial sections from tibialis anterior muscle of control (-Cre, Osmr^fl/+^) or conditional knockout (+Cre, Osmr^fl/fl^ Pax7^CreERT2^) animals following deletion and 2 rounds of cardiotoxin injury and regeneration. Results are plotted as Pax7^+^ nuclei per 100 fibers (left) or Pax7^+^ nuclei as a percentage of all visualized nuclei (right).
f. Model for OSM mediated quiesence induction during skeletal muscle regeneration. Under homeostatic conditions, stem cells (grey) are deeply quiescent but can become activated and proliferate (red) following injury. As regenerated myofibers begin to form, secretion of OSM (green) allows quiescence induction by actively promoting cell cycle exit within a subset of activated satellite cells while also maintaining stem cell identity. At later timepoints, 2-3 weeks following injury, upregulation of intrinsic regulators such as Sprouty1 matures the re-selected satellite cell pool back toward the deeply quiescent homeostatic state.

OSMR-deficient mice demonstrated no difference in stem cell number or myofiber cross-sectional area in the steady state (Supplementary Fig. 6d,e); this is in keeping with the reappearance of OSM protein during myofiber repair post injury (Fig. 4a), and consistent with a role in re-establishment of quiescence following activation. Serial injury of OSMR-deficient muscle would therefore be expected to deplete the satellite cell pool. Indeed, two rounds of cardiotoxin injury in OSMRβ-deficient mice resulted in a substantial decrease in satellite cell numbers (Supplementary Fig. 7a), associated with a marked regenerative block and >50% reduction in mean myofiber cross-sectional area (CSA; Supplementary Fig. 7b,c,d). To exclude that this reflected effects of OSM on other cell populations such as inflammatory cells, we used Cre-loxP mutagenesis to conditionally ablate OSMRβ in Pax7^+^ MuSC using inducible lineage-restricted Cre expression^35^. Serial muscle injury of conditionally deleted animals led to impaired muscle regeneration, marked decrease in myofiber cross sectional area, and severe depletion of satellite cells (Fig. 4c,d,e; Supplementary Fig. 7e). The regenerative defect of satellite-cell specific OSMR deletion was indistinguishable from that of whole-body constitutive deletion, demonstrating a specific requirement for OSM signaling in satellite cells.

## Discussion

Here we describe a novel *in vivo* imaging-based screen which identifies Oncostatin M as a key niche-derived secreted regulator of muscle stem cell quiescence. OSM is, to our knowledge, one of only two niche-localized factors known to induce quiescence in solid organ stem cells^36^, and the first shown to maintain transplantation potential. The ability of OSM to promote MuSC engraftment without inducing proliferation is distinct from previously reported factors such as regulators of p38 MAPK^6, 10^ as well as related cytokines such as IL-6 and LIF (Fig. 2h)^18, 20, 21, 37^. The quiescence-inducing function of OSM in skeletal muscle is likewise distinct from its effect in other tissues such as the heart, where it acts to induce cardiomyocyte Proliferation^33^, and more closely resembles the cytostatic effect originally described in cancer cell lines and mouse myoblasts^38, 39, 40^. This cytostatic effect was moreover shown to be associated with STAT3 activation, and inhibition of CDK2 activity. Although OSM was paradoxically reported to have a pro-proliferative effect on human myogenic progenitor cells, interpretation of this finding is complicated by the fact that human (but not mouse) OSM also signals through the proliferation-inducing LIF receptor (LIFR) pathway^41, 42^.

The temporal expression pattern of myofiber-derived OSM resembles that of Sprouty1, a satellite cell-intrinsic inhibitor of FGF signaling whose re-expression following injury is likewise critical for re-induction of stem cell quiescence^9^. Importantly, while OSM is re-expressed by fibers within 4-10 days of injury, Sprouty1 re-expression requires 3 weeks^9^. We suggest that this difference may define an early stage of quiescence, in which nascent myofibers secrete OSM in order to induce cell cycle exit and maintain stem cell identity in a subset of activated satellite cells (Fig. 4f). This model is in keeping with restricted expression of OSMR within a subset of satellite cells (Supplementary Fig. 2b), and adds to our knowledge of heterogeneity within the stem cell pool^43, 44^ In addition to its role following muscle injury, we speculate that OSM signaling may play additional roles in the steady state, for instance preventing full activation of stem cells induced by remote damage to enter the “G_Alert_” state^7^. Future work will address this possibility, the fate of OSMR deficient MuSC following activation, as well as whether dysregulation of OSM levels with aging or chronic injury may contribute to impaired maintenance and function of the stem cell pool^3, 4, 6, 28^. It will likewise be important to understand the mechanisms by which signaling pathways shared by multiple IL-6 family members are translated into distinct biological responses ^44, 45^. Finally, we anticipate that the *in vivo* imaging-based screening approach reported here will prove useful in identifying additional niche components controlling the function of satellite cells and stem cells in other lineages.

## Methods

### Mice

Experimental protocols involving mice were reviewed and approved by the Stanford University Administrative Panel on Laboratory Animal Care and the GNF Institutional Animal Care and Use Committee. All experiments were performed in compliance with institutional guidelines. All experiments utilized adult (8-12 week old) female mice. Hindlimb-irradiated NOD/SCID mice (Jackson Laboratory) were used as stem cell recipients during screening of protein library pools. Subsequent experiments involving stem cell transplants utilized hindlimb-irradiated, immunocompromised NSG mice. Ubiquitously expressing transgenic CAG-*luciferase-GFP* mice were obtained and maintained as described previously^6, 25^. Lineage reporter Pax7-IRES-GFP mice were generated using a strategy similar to one previously described^46^. In brief, to generate a targeting vector for homologous recombination, Pax7 genomic sequences were amplified from C57BL/6 mouse genomic DNA and Pax7 homology arms were cloned into a targeting vector containing IRES-EGFP and FRT-flanked neomycin cassettes. C57BL/6 mouse ES cells were transfected by electroporation of linearized Pax7 targeting vector. Transfected ES cells were selected for neomycin resistance using 0.2 mg/ml Geneticin (Invitrogen #10131-019). Ten days after transfection, 300 G418-resistant ES cell clones were isolated and genotyped by PCR for homologous recombination with primers CTATCGCCTTCTTGACGA (neomycin) and GTGTCCCTGGTGTCTTGT (Pax7). Correct targeting was confirmed by Southern blot using a neomycin-specific probe, which allowed the exclusion of random integration events of the targeting vector. Selected targeted ES cells were injected into BALB/c blastocysts and chimeric mice were bred with C57BL/6 females to obtain Pax7-IRES-GFP knock-in mice. To eliminate the FRT-flanked neomycin cassette, Pax7 gene targeted mice were crossed with a mouse line expressing Flp recombinase and analyzed for the loss of the *neomycin* cassette. Pax7-GFP mice without *neo* were used for further analysis.

The Osmr allele was obtained from Jax (B6;129-Osmrtm1.1Nat/J) and typed via real time PCR (Transnetyx). For complete deletion, floxed mice were bred to Thy1-cre to induce deletion in the germline. The deleted allele was confirmed by PCR and bred away from the Cre driver. The strain was maintained by intercross of heterozygous deleted animals. For conditional deletion, the floxed allele was bred to Pax7^CreERT2 35^. To induce deletion, Osmr^fl/fl^ Pax7^CreERT2^ mice or controls (floxed homozygous or heterozygous without Cre driver) were injected on five consecutive days with 200 ul of 20 mg/ml tamoxifen in corn oil (Sigma), and then subsequently maintained on tamoxifen-containing chow (TD.130855, Harlan-Teklad) in order to prevent counterselection against deleted satellite cells^47^.

### Stem Cell Purification and Culture

Mouse muscle stem cells were purified essentially as previously described^6^. In brief, tibialis anterior and gastrocnemius muscles were dissected and digested with Collagenase II and Dispase (Worthington Biochemical Corporation) for a total of 90-120 minutes. Following centrifugation and resuspension, the cell suspension was strained a 40-μm filter (BD Biosciences). Cells were stained with the following biotinylated antibodies: anti-CD45 (BD Biosciences, clone 30F11, catalog # 553078, 1:500), anti-CD11b (BD Biosciences, clone M1/70, catalog # 553309, 1:800), anti-CD31 (eBiosciences, clone 390, catalog # 13-0311-82, 1:200) and anti-Sca1 (BD Biosciences, clone E13-161.7, catalog # 553334, 1:200). Cells were then incubated with streptavidin (SA) microbeads (Miltenyi Biotech), SA-Texas Red (Invitrogen, catalog # S872, 1:200), PE labeled anti-integrin α7 antibody labeled with phycoerythrin (AbLab, clone R2F2, catalog # 10ST215, 1:500) and anti-CD34 eFluor660 (eBioscience, clone RAM34, catalog # 50-0341-82, 1:67). In some experiments, cells were stained with eFluor 450 directly conjugated lineage antibodies (anti-CD45, anti-CD31, anti-Sca1, anti-CD11b, anti-Ter119; eBioscience). Dead cells were stained with propidium iodide. Cells sorting on a FACS Aria 2 routinely yielded >95% CD34^+^α7 integrin^+^ cells from a single round of sorting. Scatter plots were analyzed using FlowJo (TreeStar). All *in vitro* culture experiments were performed using media changes with fresh recombinant protein added every 3 days. For screening, His-tagged cDNA clones comprising the GNF Secretomics Collection were expressed in 293T Freestyle cells and affinity purified using nickel agarose using in-house automation instrumentation. Retained proteins were eluted in TBS with 250 mM imidazole and were pooled manually. Pooled proteins were added to 800 purified stem cell counts at 10% final culture volume; no attempts were made to normalize the concentration of individual proteins within the pools, which therefore varied as a function of protein expression and elution efficiency. Media was changed after 3 days and fresh protein-containing media was added for another 3 days prior to transplant. MuSC were cultured under minimal conditions on plastic without exogenous extracellular matrix, conditions that favor commitment and differentiation ^48^. Standard muscle stem cell culture was performed in DMEM/F10 (50:50), 15% FBS, 1% penicillin-streptomycin, and 2.5 ng/ml rh bFGF. For screening experiments, culture was performed in medium above but without FGF. Pilot experiments indicated that use of collagen coated plates dramatically reduced assay window, therefore uncoated plastic plates were used throughout, including for long-term (14 day) stem cell culture. Recombinant Oncostatin M (R&D Systems), recombinant human basic FGF (Promega), SB202190 (EMD Chemicals) and U0126 (Cell Signaling) were obtained commercially. Surface area for experiments performed in chamber slides (Figure 2A) was 0.8 cm^2^, with an initial seeding density of ~31 cells/mm^2^.

### Muscle stem cell transplantation and muscle injury

Stem cell harvesting, intramuscular injection, and cardiotoxin-mediated muscle injury were all performed essentially as previously described^6^, with the exception that 50 ul 10 μM cardiotoxin was used for all injuries. For counted cell transplants, stem cells were counted manually using a hemocytometer prior to transplant.

### Bioluminescence imaging

Bioluminescence imaging (BLI) was performed using either a Xenogen 100 or IVIS Spectrum CT, as previously described^6, 25, 48^. Briefly, 150 μL D-luciferin (reconstituted at 30 mg/mL in sterile water; Caliper LifeSciences) was administered by intraperitoneal injection into anesthetized mice. BLI were acquired using 60 second exposures for a total of 12 min following luciferin injection. Images were recorded and analyzed using Living Image software (Caliper LifeSciences). For images acquired on the Xenogen 100, a signal value of 100,000 photons per second was routinely used to define the engraftment threshold, as previously described^6, 25^. Engraftment thresholds were approximately ten-fold higher for imaging performed on the IVIS Spectrum CT, reflecting the higher sensitivity on this instrument and intrinsic detector characteristics between machines. *In vitro* imaging was performed similarly but with a 1:1000 final dilution of stock luciferin.

### Graphical and Statistical Analysis

Graphs and statistics were generated using GraphPad Prism software. Student’s two-tailed t tests were used to calculate p values.

### Cell and tissue staining

Tibialis anterior (TA) muscle sections were prepared for staining essentially as previously described^6, 48^. For OSMR staining, whole TA muscles were first fixed with 0.5% paraformaldehyde/PBS for 2 hr at 4C, followed by incubation in 20% sucrose/PBS overnight at 4C. 10 μM sections were cut for all stainings. Antibodies were: anti-laminin (Millipore, catalog #05-206, 1:250), anti-GFP (Invitrogen, catalog #A11122, 1:200), anti-OSMR (R&D, catalog #AF665, 1:200), anti-Pax7 (Santa Cruz Biotechnology, catalog #sc81648, 1:50; or Developmental Studies Hybridoma Bank, 2 μg/mL final concentration), AlexaFluor 594-conjugated donkey anti-rat IgG1 and AlexaFluor 488-conjugated donkey anti-rabbit (Jackson ImmunoResearch, catalog # 712-585-150 and 711-545-152 respectively, 1:200 each). Nuclei were counterstained with either DAPI (Invitrogen) or TO-PRO-3 (Invitrogen). Images were acquired with an AxioPlan2 epifluorescent microscope (Carl Zeiss) with ORCA-ER digital camera (Hamamatsu Photonics).

### Single-cell RT-PCR

Single cell RT-PCR was performed as described^6, 25^. In brief, MuSCs were obtained from digested muscle as above, and three rounds of FACS sorting was performed in order to obtain a highly purified CD34^+^α7 integrin^+^ MuSC population. In the last sorting step, the cells were dispensed into PCR tubes containing lysis buffer under single cell conditions (1 cell/well) or non-single cell conditions (50 cells/well). Multiplexed RT-PCR was performed using the following primers: Pax7 ext FWD GAGTTCGATTAGCCGAGTGC; Pax7 ext REV GGTTAGCTCCTGCCTGCTTA; Pax7 int FWD GCGAGAAGAAAGCCAAACAC; Pax7 int REV GGGTGTAGATGTCC GGGTAG Myf5 ext FWD AGACGCCTGAAGAAGGTCAA; Myf5 ext REV AGCTGGACACGGAGCTTTTA; Myf5 int FWD C CACCAACC CTAAC CAGAGA; Myf5 int REV CTGTTCTTTCGGGACCAG AC; Myod1 ext FWD TACCCAAGGTGGAGATCCTG; Myod1 ext REV GTGGAGATGCGCTCCACTAT; Myod1 int FWD GCC TTCTACGCACCTGGAC; Myod1 int REV ACTCTTCCCTGCCTGGACT

### EdU labeling

For *in vitro* labeling experiments, satellite cells were treated with 10 μM 5-ethynyl-2′-deoxyuridine for the last 12 hours of culture. Cells were fixed, permeabilized, and EdU incorporation detected using the Click-iT EdU Alexa Fluor 488 Imaging Kit (Life Technologies) per manufacturer’s protocol.

### qRT-PCR

Analysis of purified cultured satellite cells was performed as described^6, 25^. RNA was isolated using the RNeasy Micro Kit (Qiagen), and reverse transcribed using SuperScript III. RT-PCR was performed using a SYBR Green PCR Master Mix (Applied Biosystems). Quantification was performed using the 2–ΔΔCt method and normalized to Gapdh. Primer sequences were as follows: p27/Kip1 REV CCGGGCCGAAGAGATTTCTG; OSMR FWD CATCCCGAAGCGAAGTCTTGG; OSMR REV GGCTGGGACAGTCCATTCTAAA; GAPDH FWD AGGTCGGTGTGAACGGATTTG; GAPDH REV TGTAGACCATGTAGTTGAGGTCA; Pax7 FWD TCTCCAAGATTCTGTGCCGAT; Pax7 REV CGGGGTTCTCTCTCTTATACTCC.

### ChIP-qPCR

Stat3-ChIP was performed following the standard protocol from the ChIP-IT Express Enzymatic kit (Active Motif, California) on chromatin isolated from primary myogenic progenitor cells (8 × 10^6^) cultured with the same culture media and condition as described for muscle stem cells. Briefly, cells were crosslinked for 10 min with 1% formaldehyde and quenched with 12.5 mM glycine for 5 min. Nuclei were obtained by incubating with 1ml hypertonic lysis buffer and protease and phosphatase inhibitors as supplied for 30 min 4 °C and releasing with 20 strokes through a dounce homogenizer on ice. Chromatin was obtained with the nuclear digestion buffer and fragmented to 300-100 base-pairs with the enzymatic shearing cocktail in 50% glycerol as provided by the kit for 12 minutes at 37°C. 10% of the sheared chromatin was saved as an input control, and 10 ug of sheared chromatin from each condition were immunoprecipitated with antibody to mouse STAT3 (610190, BD Bioscience) or normal mouse IgG (Cell Signaling) overnight at 4°C, followed by 1 h of incubation at 4 °C with Protein G-coated magnetic beads (Active Motif). Washing and reverse crosslinking was performed according to manufacturer’s protocol. The precipitated DNA was amplified and analyzed by real-time SYBR Green PCR (Applied Biosystem) on an ABI PRISM 7900 HT Sequence Detection system (Applied Biosystem). Primer pairs (EpiTect GPM105088(+)01A, SA Bioscience) designed for quantitative RT-PCR analysis were used to amplify the putative Stat3 binding site adjacent to the mouse Pax7 locus (Chr4: 139387970-139388390), identified via Biobase Transfac analysis (Biobase Corp. MA). The Stat3 occupancy was calculated as a ratio of the amplification efficiency of Stat3 ChIP over input chromatin.

### Intracellular FACS

Cells were fixed with 4% formaldehyde for 10 min at RT and permeabilized by adding cell suspension drop-wise into ice-cold 100% methanol while gentle vortexing to a final concentration of 90% methanol and incubate on ice for 30 min. Permeabilized cells were resuspended in FACS buffer (1xPBS, 1% BSA), stained with antibody for 30 min at room temperature and analyzed via FACS Calibur flow cytometer (BD, Bioscience). Phospho-Histone H3 (ser10) was performed using the Alexa Fluor 488 conjugated H3S10 primary antibody (9708, Cell Signaling). Phospho-Stat3 and Phospho-Stat5 were performed using PE-mouse anti-Stat3 (Y705) (612569, BD Bioscience) and PerCP-Cy5.5 mouse anti-Stat5 (pY694) (560118, BD Bioscience) respectively.

### Microarray analysis

Total RNA was prepared with Qiagen RNeasy kits. Twenty nanograms of total RNA was prepared for array analysis using Affymetrix 3’ IVT Express amplification reagents. Fifteen micrograms of cRNA was hybridized to Affymetrix MOE_430_2 arrays, and arrays were washed, stained, and scanned using standard methods. CEL files were processed using MAS5. Microarray data have been submitted to GEO (accession number GSE69976).

### Bioinformatic analysis

OSM signature genes were defined as the top-quartile of the most up-regulated genes using the ratio of OSM-treatment/no-OSM treatment. Similarly a quiescence signature was defined using published data^23, 24^ as the intersection of the top quartile of up-regulated genes (ratio of quiescence vs activated) in two comparisons. These two signature sets were then compared to each other and the number of overlapping genes was found to be higher than expected by chance as assessed by Fisher’s Exact Test (n=754 genes, 2.01 fold enrichment, p=4e-105).

## Data availability

Microarray data are available via GEO (accession number GSE69976). The data that support the findings of this study are available on request from the corresponding authors (SCS, HMB).

## Author Contributions

Experiments were designed, carried out, and interpreted by S.C.S., S.C.S., A.T.V.H., S.Y.C., J.M., C.S. and H.M.B. J.L. and J.W. contributed to microarray experiments and bioinformatics analysis. Pax7-GFP mice were developed by B.K. All authors contributed to writing and revision of the manuscript.

## Acknowledgements

This report is dedicated to the memory of Matt Furia. The authors thank Penney Gilbert and Ben Cosgrove for advice and methodological input; John Joslin, Sarah Cox, Mark Knuth, Tony Orth, Heath Klock, and Scott Lesley for the Secretomics library; Angela Christiano for the OSM staining protocol; Mary Frazer, Richard Eddins, Kassie Koleckar, and Peggy Kraft for animal husbandry support; and Richard Glynne for advice and support. The Stanford Center for Innovation in In-Vivo Imaging (SCI3), the Stanford Shared FACS Facility (SSFF), the Stanford Lokey Stem Cell Research FACS Facility, and the Stanford Veterinary Service Center (VSC) are thanked for technical support. This study was supported by the Baxter Foundation, NIH R01 AG020961, NIH R01 NS089533 and CIRM R5-07469 (to H.M.B.).

---

**Supplementary Figure 1.**
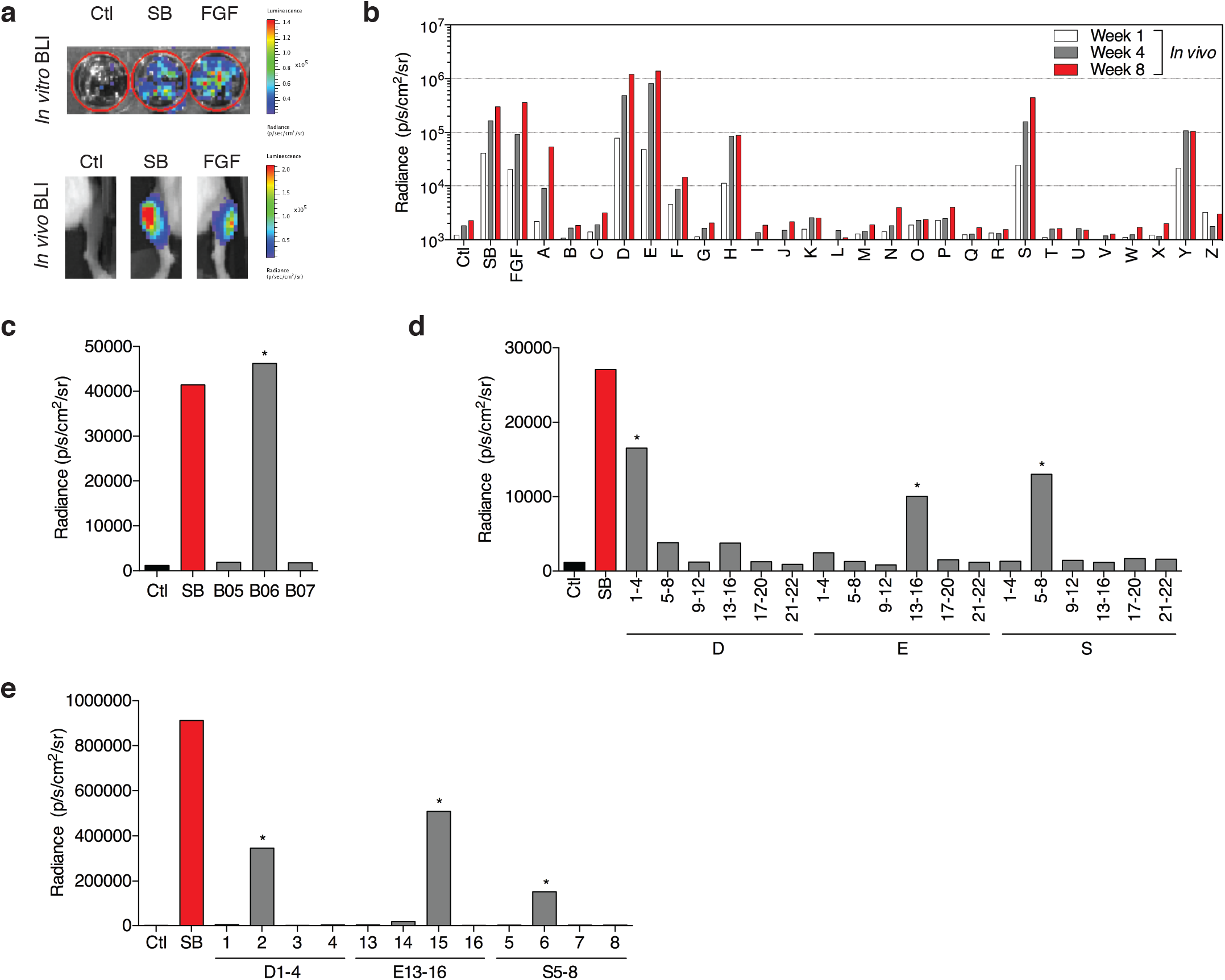
Deconvolution of screening hits. a. Representative *in* vitro and *in vivo* BLI results for the screening strategy. 800 FACS-sorted MuSC were cultured on plastic in minimal conditions either without additional factors (Ctl), with SB202190 (SB; 10 μm), or with recombinant human Fibroblast Growth Factor (FGF; 2.5 ng/mL). *In vitro* BLI was performed after 6 days, the cell were then harvested and then transferred intramuscularly. *In vivo* BLI was performed 2 weeks later.
b. Serial *in vivo* BLI from multiplexed protein screening. BLI was performed at 1 (white), 4 (grey), and 8 (red) weeks post transplant. Ctl: control buffer; SB: SB202190; FGF: recombinant human basic Fibroblast Growth Factor.
c. *In vivo* BLI data from final round of deconvolution of pool Y (see Fig. 1c). SB202190 (SB; 10 μm), and buffer (Ctl) were used as controls. Asterisk indicates final positive pool, containing the extracellular domain of TNFRSF1A.
d. *In vivo* BLI data from deconvolution of three positive pools from Fig. 1c, each containing 4 proteins. SB202190 (SB; 10 μm), and buffer (Ctl) were used as controls. Imaging was performed at day 10 following intramuscular transplant. Asterisk indicates positive subpools.
e. *In vivo* BLI data from final round of deconvolution of positive subpools indicated in (c). Note that each subpool now contains an individual protein. Imaging was performed at day 14 following intramuscular transplant. Asterisk indicates positive subpools, each containing Oncostatin M.

**Supplementary Figure 2.**
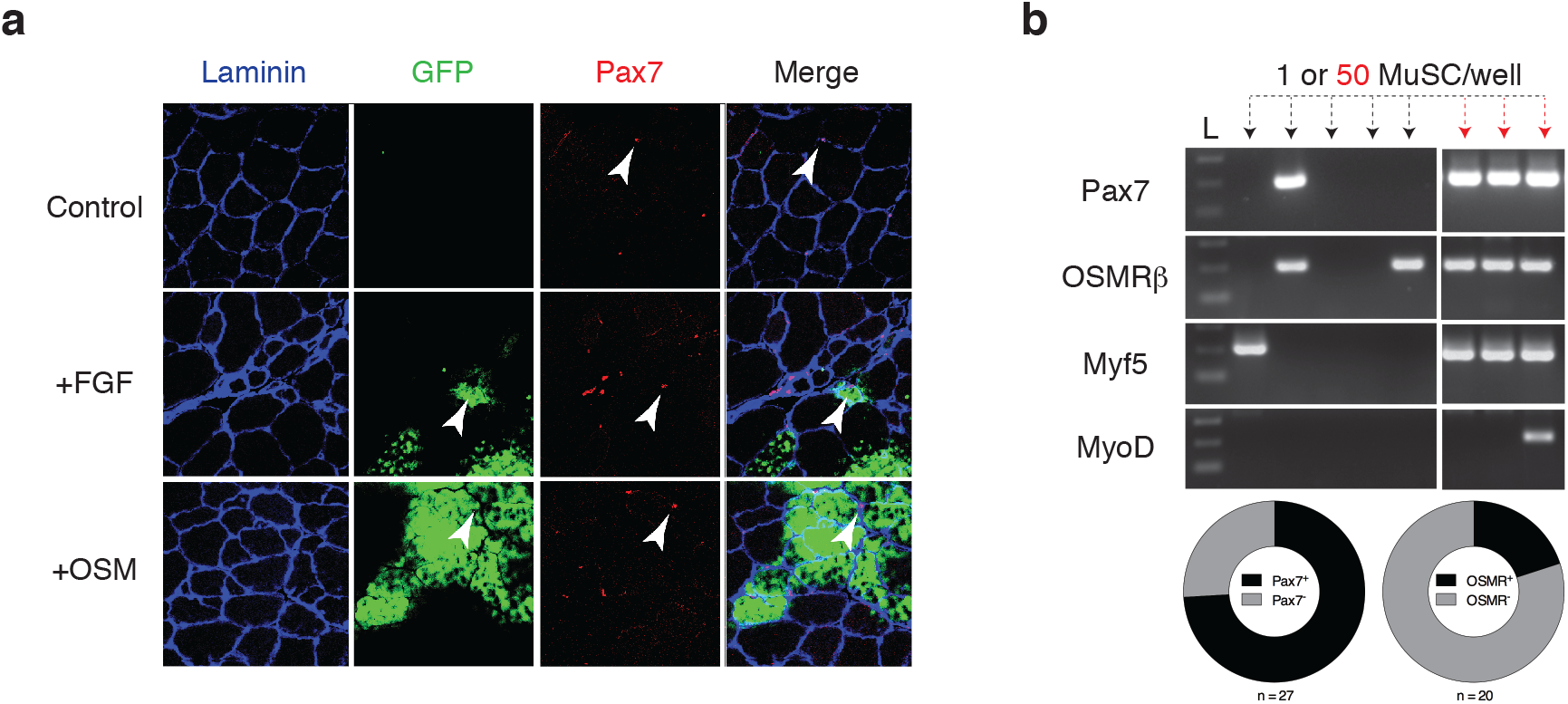
Expression of OSMRβ within muscle stem cells. a. Representative immunofluorescence analysis of muscle sections following transplant with either buffer (top row), FGF (middle row) or OSM (bottom row) treated stem cells isolated from CAG-*luciferase-GFP* mice. Note that engrafted fibers derived from these cells express both luciferase as well as GFP. White arrowheads indicate Pax7^+^ satellite cells adjacent to areas of GFP^+^ donor-derived fibers.
b. OSMRβ expression on single sorted MuSC. Triple FACS-sorted MuSCs were sorted into PCR tubes at either 1 cell per well or 50 cells per well (positive control) for RT-PCR. Representative data from nested, non-quantitative RT-PCR demonstrating expression of Pax7, OSMRβ, Myf5, and MyoD in either single cells (black arrows) or in pools of 50 cells (red arrows). Of cells demonstrating amplification, OSM^+^/Pax7^+^ was ~19% (20 total), OSM^+^Myf5^+^/Myf5^+^ was ~13% (13 total), Pax7^+^ was 75% (27 total), Myf5^+^ was ~48% (13 total). Schematic at bottom summarizes Pax7 and OSMRβ results.

**Supplementary Figure 3.**
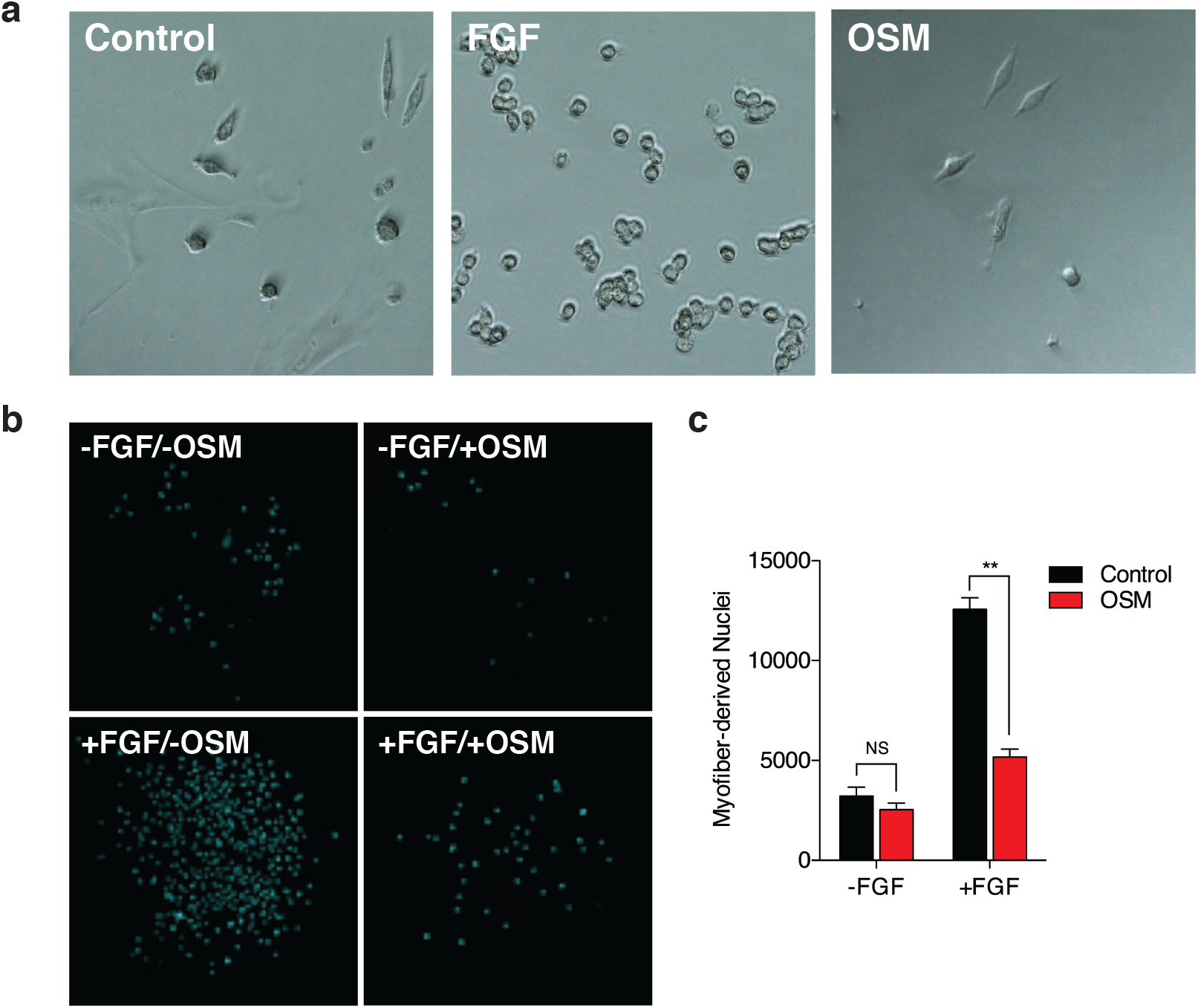
OSM represses proliferation of isolated and fiber-associated muscle stem cells. a. Phase contrast micrograph images of FACS-sorted MuSC culture for 6 days with either control buffer (left), FGF (2.5 ng/ml; middle), or OSM (100 ng/ml; right). Images were acquired at 20X.
b. Imaging of cells derived from myofiber cultures described in Fig. 3c. Bulk muscle fibers derived from C57BL/6 mice were cultured for 5 days in the simultaneous absence or presence of FGF and/or OSM. Plate-bound cells were quantified by DAPI staining, high-content imaging, and automated counting of cell nuclei.
c. Quantification of myofiber-derived colony formation. Bulk muscle fibers from wild type C57BL/6 mice were cultured for 5 days in the absence or presence of FGF and OSM. At the end of the culture period, plate-bound cells representing progeny of activated satellite cells were quantified by high-content imaging and automated counting of DAPI^+^ nuclei (see Supplementary Fig. 3b). NS, not significant; double asterisk indicates p<0.01.

**Supplementary Figure 4.**
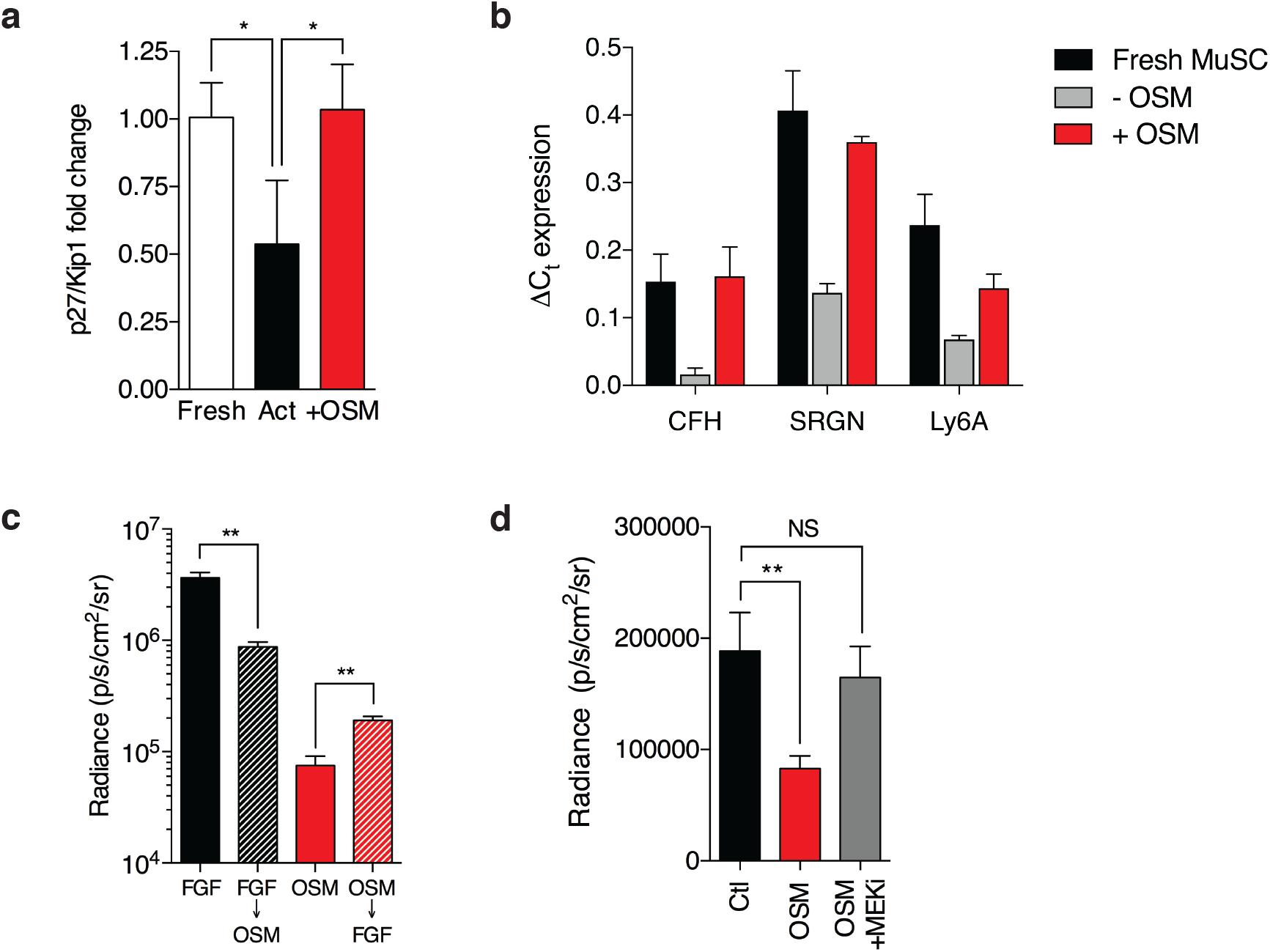
Mechanism of OSM-induced quiescence. a. p27/Kip1 induction in response to OSM treatment *in vitro*. qPCR was performed on FACS-sorted MuSC either freshly isolated (Fresh) or cultured for 6d in the absence (Activated, Act) or presence (+OSM) of OSM. Data were normalized to expression of GAPDH. Mean +/- SD is shown, n=3 replicates; asterisk indicates p<0.05.
b. Validation of quiescence gene signature. Expression of three genes from the microarray-derived quiescence signature was independently assessed by qPCR analysis of fresh MuSC or MuSC cultured without (-OSM) or with OSM (+OSM).
c. *In vitro* BLI for FACS-sorted MuSC from Fig. 2e. Cells were cultured for 6 days under the indicated conditions, and BLI was performed prior to cell fixation. Mean +/- SD is shown, n=4 replicates.
d. Proliferation of FACS-sorted MuSC cultured for 6 days in the presence or absence of OSM, or with both OSM and the MEK inhibitor U0126 (5 μM). Mean +/- SD is shown, n=3 replicates.

**Supplementary Figure 5.**
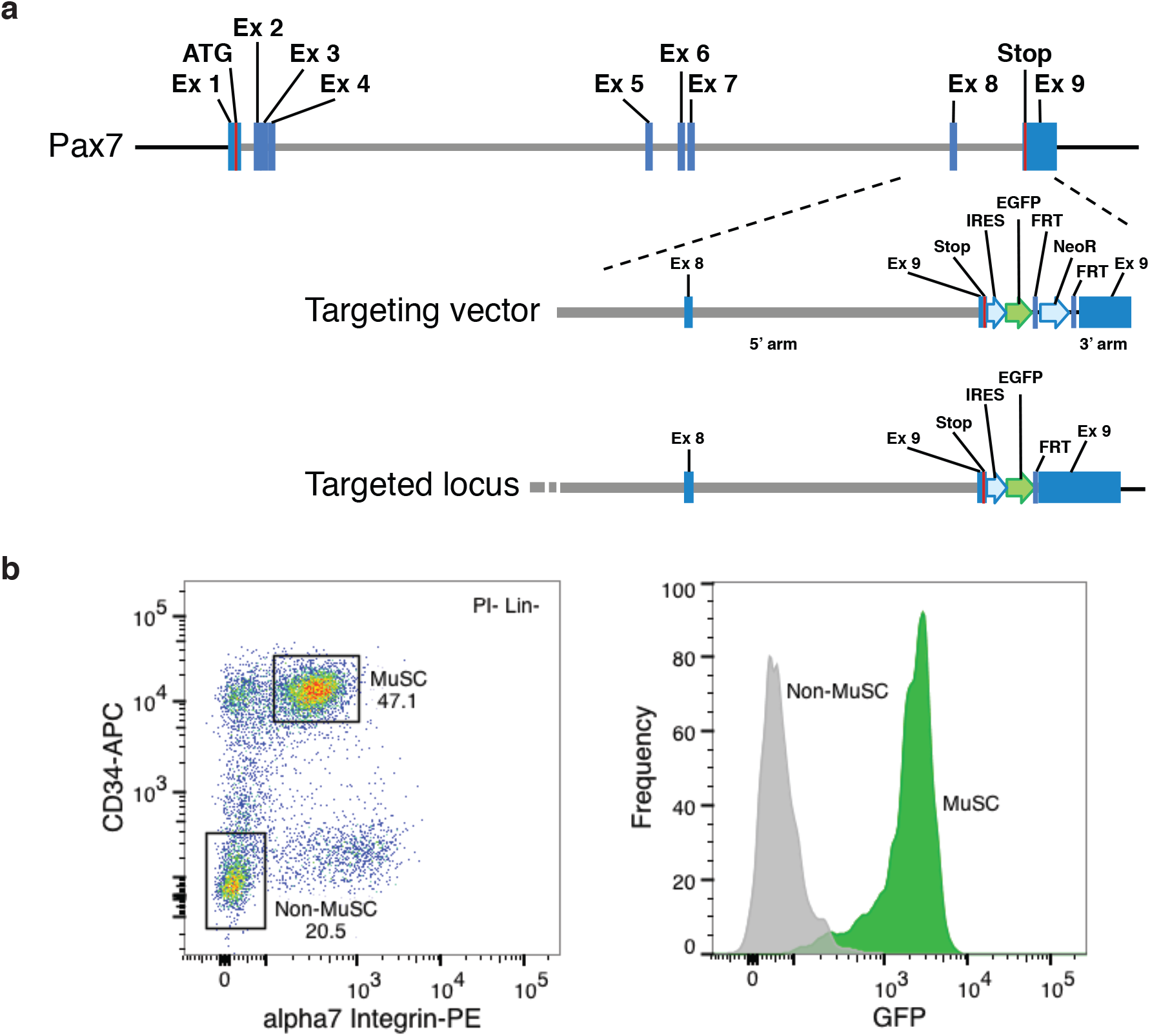
Generation and validation of Pax7^GFP^ allele. a. Schematic of targeting vector used to generate Pax7^GFP^ mice, containing insertion of an Internal Ribosome Entry Site and Green Fluorescent Protein (IRES-GFP) cassette into the 3’ untranslated region of the *Pax7* locus.
b. FACS analysis of the muscle-derived mononuclear fraction from homozygous Pax7^GFP^ mice. Left: Gating of PI^−^CD11b^−^Scal^−^CD31^−^CD45 cells into MuSC and non-MuSC populations by CD34 and α7-Integrin expression. Right: GFP expression in MuSC (PI^−^CD11b^−^Sca1^−^CD31^−^CD45^−^CD34^+^α7 integrin^+^; green) and non-MuSC (PI^−^CDllb^−^Scal^−^CD31^−^CD45^−^CD34^+^α7 integrin^−^; grey) populations.

**Supplementary Figure 6.**
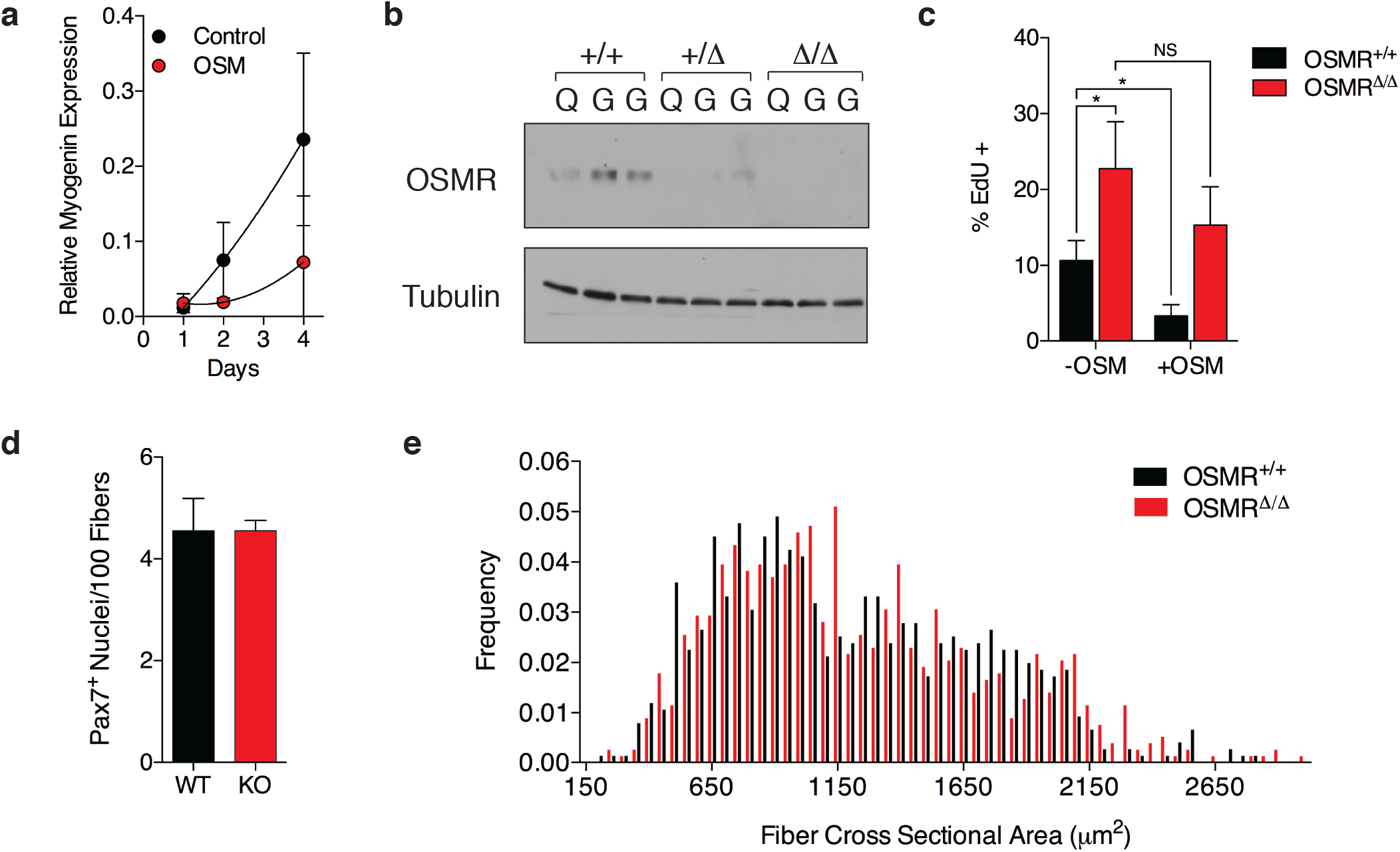
OSMR is required for OSM signaling in muscle stem cells. a. qRT-PCR analysis of *myogenin* expression following 6 day culture of FACS-sorted MuSC in the presence of OSM (gray circles) or buffer control (black). Mean +/- SD is shown, n=3 replicates.
b. Western blot analysis of uninjured quadriceps (Q) or gastrocnemius (G) muscle from wild type (+/+), heterozygous (Δ/+) or homozygous (Δ/Δ) mutant OSMRβ animals.
c. Increased proliferation of OSMRβ-deficient MuSC. FACS-sorted wild type (OSMR^+/+^) or homozygous OSMRβ mutant (OSMR^Δ/Δ^) muscle stem cells were cultured for 6 days without (-OSM) or with OSM (+OSM). Cells were pulsed labeled with EdU for the final 12 hours of culture.
d. Quantitation of Pax7+ stem cells in serial sections from tibialis anterior muscle of uninjured wild type (+/+) or homozygous (Δ/Δ) mutant OSMRβ animals. Results are expressed relative to fiber number.
e. Cross-sectional area (CSA) distribution of myofibers from tibialis anterior (TA) muscle of uninjured wild type (+/+; open boxes) or homozygous (Δ/Δ; black boxes) mutant OSMRβ animals. >750 fibers were quantified per genotype.

**Supplementary Figure 7.**
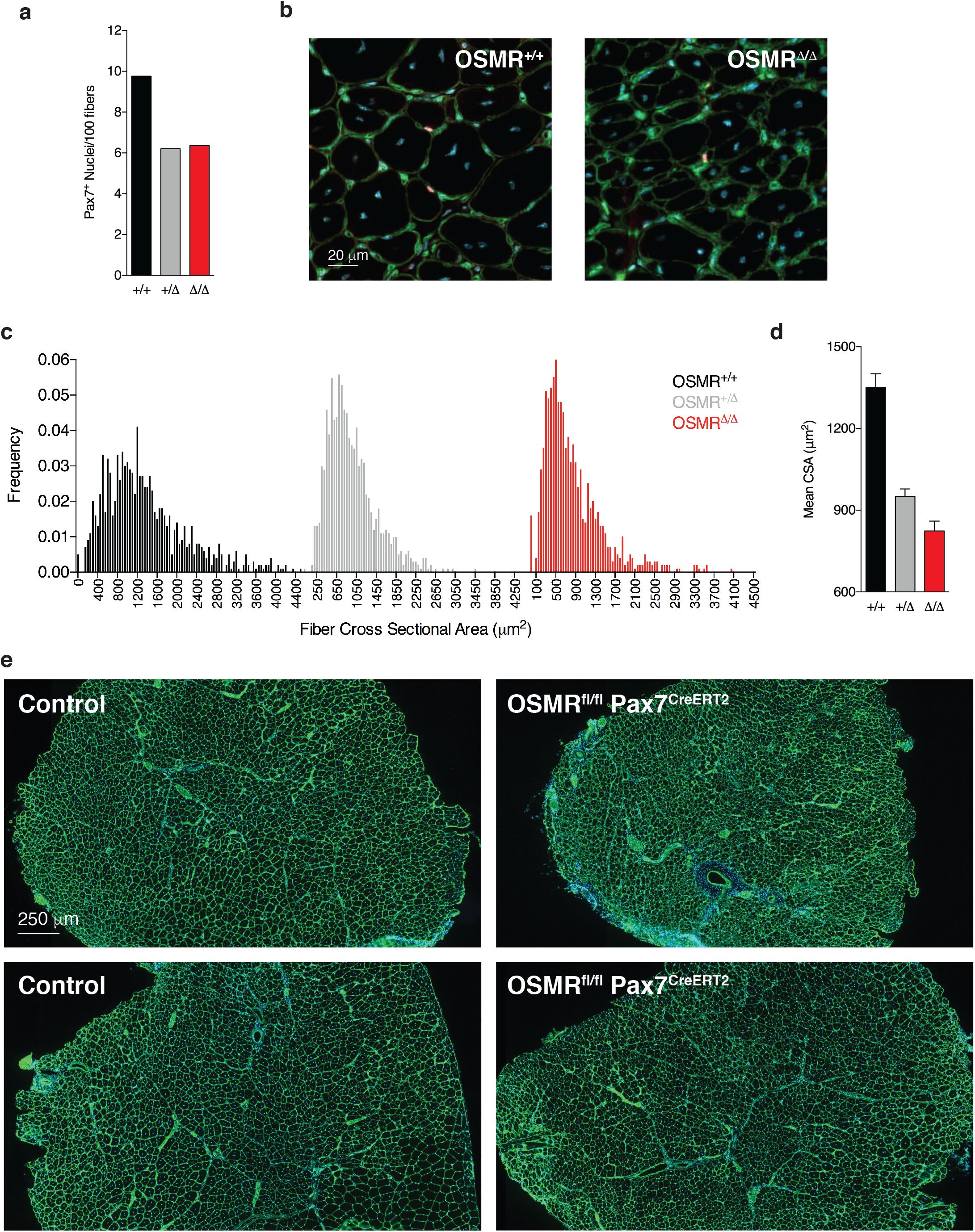
Characterization of muscle regeneration in OSMRβ deficient mice. a. Quantitation of Pax7^+^ stem cells in serial sections from tibialis anterior muscle of wild type (+/+), heterozygous (Δ/+) or homozygous (Δ/Δ) mutant OSMRβ animals following 2 rounds of cardiotoxin injury and regeneration. Results are expressed relative to fiber number.
b. Representative immunofluorescence images from tibialis anterior muscle of wild type (+/+) or homozygous mutant (Δ/Δ) OSMRβ animals following 2 rounds of cardiotoxin injury and regeneration. Blue: DAPI; Green: Laminin; Red: Pax7.
c. Cross-sectional area (CSA) distribution of myofibers from tibialis anterior (TA) muscle of wild type (+/+; open boxes), heterozygous (Δ/+; gray boxes) or homozygous (Δ/Δ; black boxes) mutant OSMRβ animals following 2 rounds of cardiotoxin injury and regeneration. >1000 fibers were quantified per pooled genotype.
d. Quantification of mean CSA from TA muscle section described in (e). >1000 fibers were quantified per pooled genotype. 95% confidence intervals of the mean are indicated.
e. Low magnification images of immunofluorescence images from tibialis anterior muscle of control (Osmr^fl/+^) or conditional knockout (Osmr^fl/fl^ Pax7^CreERT2^) animals following deletion and 2 rounds of cardiotoxin injury and regeneration. Blue: DAPI; Green: Laminin.

